# Joint profiling of 5mC, 5hmC, and the transcriptome in single cells identifies factors responsible for genome-wide DNA methylation erasure in human primordial germ cell maturation

**DOI:** 10.1101/2025.04.01.646736

**Authors:** Alex Chialastri, Kellie A. Heom, Joanna J. Gell, Fei-man Hsu, Sissy E. Wamaitha, Amander T. Clark, Siddharth S. Dey

## Abstract

Reprogramming of DNA methylation plays a vital role in the establishment of cell identity during early mammalian development. To gain deeper mechanistic insights into this process requires capturing the entire dynamics of DNA methylation – both 5-methylcytosine (5mC) and its downstream oxidation product 5-hydroxymethylcytosine (5hmC) – in individual cells. Therefore, in this work, we report a new single-cell genome-wide strand-specific sequencing method, scMHT-seq, to jointly profile 5mC, 5hmC, and the transcriptome from individual cells. Using human embryonic stem cells (hESCs), we first show that scMHT-seq can accurately detect both 5mC and 5hmC from the same cell with minimal crosstalk in quantifying these two DNA modifications, and that the multi-modal measurements are in close agreement with individual measurements of 5mC and 5hmC in single cells. After establishing the method, we next applied scMHT-seq to gain insights into human primordial germ cell (hPGC) development. After specification, hPGCs undergo rapid global demethylation as they mature, and this reprogramming is critical for normal development of gametes. However, it has not been possible to fully overcome this key epigenetic barrier in culture, thereby limiting our ability to generate mature hPGC-like cells (hPGCLCs) and accomplish *in vitro* gametogenesis. To gain deeper understanding of the molecular factors involved in germ cell maturation, we applied scMHT-seq to an extended *in vitro* culture system for generating hPGCLCs and observed partial and heterogeneous erasure of the methylome across single cells that is mechanistically predominantly driven by passive demethylation due to reduced DNMT1-mediated maintenance methylation activity. Notably, we discover that hPGCLCs in extended culture can be transcriptionally classified into two distinct states, with one population enriched with more mature hPGCLCs exhibiting genome-wide loss of DNA methylation. Moreover, analysis of these two cell states identifies DND1 and SOX15 as two factors that are potentially key drivers of hPGCLC demethylation and maturation. Overall, we demonstrate that scMHT-seq is a robust and high-throughput technology that can provide insights into the mechanisms driving DNA methylation dynamics and their effect on cell states.

## INTRODUCTION

DNA methylation (5-methylcytosine or 5mC) is a well-established regulator of gene expression that plays a key role in impacting cellular identity^1^. In this context, during early development in mammals, DNA methylation erasure is central in generating pluripotent epiblast cells prior to implantation, and again in the development of the germline to reset the epigenome of the gametes^1,2^. For example, during the maturation of human primordial germ cells (hPGCs), genome-wide DNA methylation levels drop from over 80% to 5%^3^, with this demethylation having key roles in X chromosome reactivation in females, and in removing 5mC from imprinted and meiotic genes to erase its prior cell identity and to facilitate differentiation into mature gametes^2^. While the global loss of DNA methylation is a key epigenetic barrier necessary for hPGC maturation, the mechanisms driving 5mC removal during hPGC development are not well understood, unlike in mouse PGCs, where both passive demethylation – arising from a lack of DNMT1-mediated maintenance methylation – and active demethylation – arising from the conversion of 5mC to 5hmC due to the activity of TET proteins – have been observed (Fig. 1a)^4–7^. Gaining deeper insights into the erasure of DNA methylation is central to mimicking the maturation of hPGC-like cells (hPGCLC) in a dish. Recently, several *in vitro* models of human germ cell development have been established, in which human embryonic stem cells or induced pluripotent stem cells are differentiated into hPGCLCs that express key early hPGC markers such as SOX17, PRDM1, and TFAP2C; however, these hPGCLCs exhibit limited global DNA demethylation, a key bottleneck in accomplishing *in vitro* gametogenesis^2,8–10^. Therefore, in this work, we employ a long-term hPGCLC culture system we developed previously to investigate the role of DNA methylation in germ cell development *in vitro* and to identify molecular factors involved in the erasure of the methylome and maturation of hPGCLCs^11^. More generally, given the importance of DNA methylation in controlling cellular memory and phenotypes, it is critical to map the dynamics of this modification as well as infer the mechanisms driving reprogramming of the methylome to gain deeper insights into cell state transitions. To fully characterize the turnover associated with the gain and loss of DNA methylation and to directly understand its relationship to cell states, it is critical to jointly quantify DNA methylation, as well as its downstream oxidation product DNA hydroxymethylation, at a single-nucleotide and strand-specific resolution, together with mRNA from single cells. However, accomplishing all these three measurements from single cells has not been reported previously; therefore, in this work we have developed a new method scMHT-seq to simultaneously profile DNA <u>m</u>ethylation, DNA <u>h</u>ydroxymethylation and the transcriptome from individual cells.

**Figure 1.**
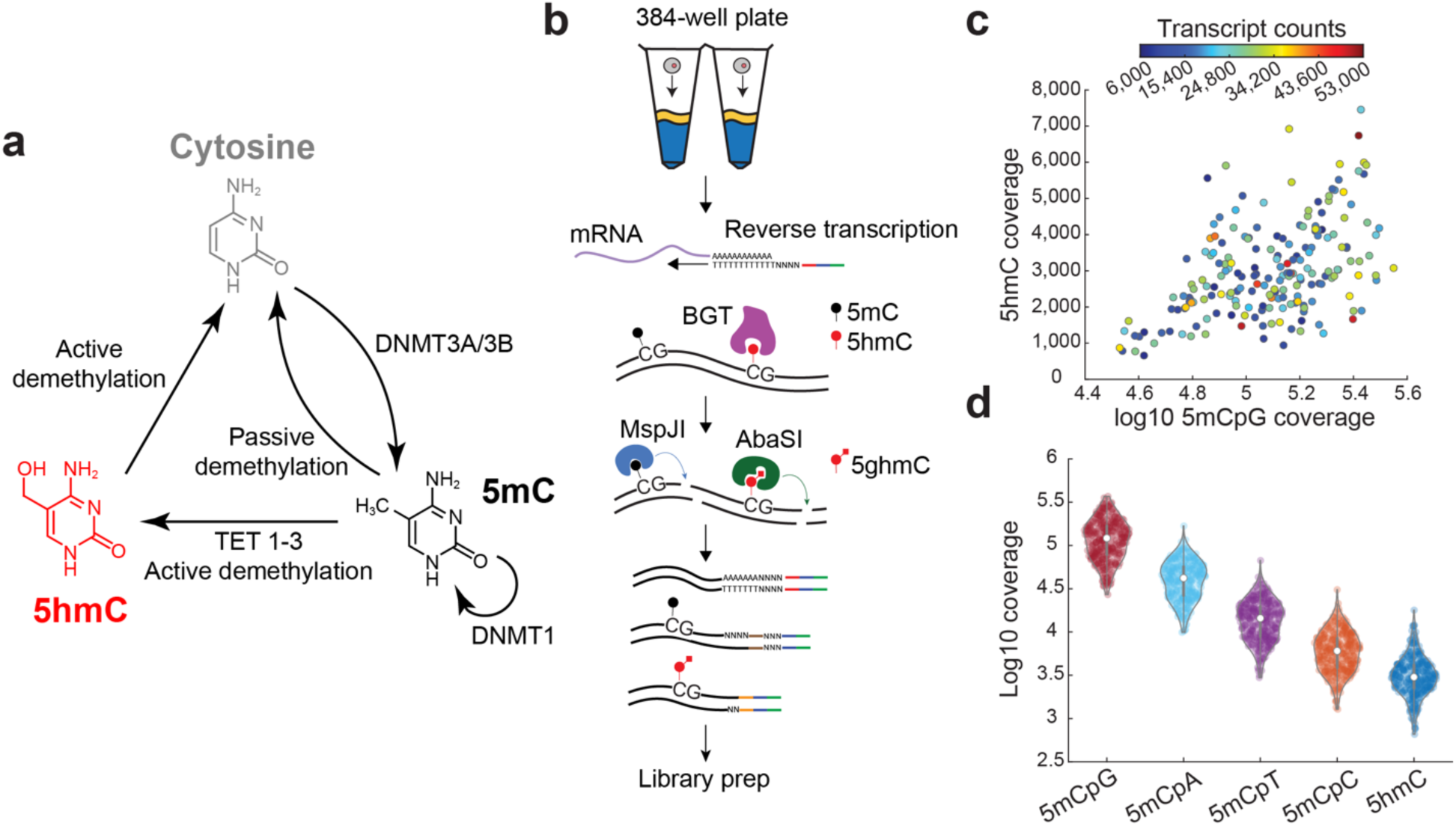
scMHT-seq enables joint profiling of DNA methylation, DNA hydroxymethylation, and the transcriptome in single cells. (**a**) Dynamics of cytosine methylation in mammalian cells. Unmethylated cytosine is converted to 5mC by the action of the *de novo* methyltransferases DNMT3A and DNMT3B, and methylation maintenance, the copying of 5mC from the old to the new DNA strand, is mediated by DNMT1. Demethylation can occur ‘passively’ through replicative dilution arising from impaired DNMT1 activity or ‘actively’ through conversion to 5hmC and other downstream oxidation products, which are not maintained through cell division and can also be enzymatically transformed, ultimately getting converted back to unmethylated cytosine. (**b**) Schematic of scMHT-seq shows that 5mC, 5hmC and mRNA can be simultaneously quantified from the same cell without requiring physical separation of the nucleic acids prior to amplification. Cell- and molecule-of-origin-specific barcodes are shown in red, brown, and gold for mRNA, 5mC, and 5hmC, respectively. The Illumina read 1 sequencing primer is shown in blue, and the T7 promoter is shown in green. (**c**) Unique 5mCpG sites, 5hmC sites, and mRNA transcripts detected using scMHT-seq in individual H9 human embryonic stem cells. (**d**) Unique 5mCpG, non-CpG methylation, and 5hmC sites detected by scMHT-seq in individual H9 cells.

Over the last few years, several groups, including ours, have developed methods to profile DNA methylation or DNA hydroxymethylation in single cells^12–16^. However, to completely elucidate the methylome dynamics, it is important to quantify both DNA methylation and DNA hydroxymethylation from single cells. Therefore, more recently, methods to jointly profile 5mC and 5hmC have been developed. For example, one approach employs a DNA hairpin to capture and denature endogenous DNA, followed by the synthesis of a copy strand that then leverages differential chemical reactivity between the methylated or hydroxymethylated endogenous DNA and the unmodified copy strand, thereby encoding the methylation status in an aligned pair of bases that can be resolved as one of six DNA states, including the 4 nucleic acid bases, 5mC and 5hmC^17^. Similarly, we developed a method that uses a combination of restriction enzymes and nucleobase conversion reactions to profile 5mC and 5hmC from the same sample at the resolution of individual CpG dinucleotides^18^. However, these methods are limited to profiling bulk samples and therefore cannot capture heterogeneity in DNA methylation dynamics between individual cells. To overcome this limitation, single-cell methods to jointly quantify 5mC and 5hmC have been developed. For example, Joint-snhmC-seq enables these combinatorial measurements by splitting genomic DNA molecules into separate reaction wells for 5mC and 5hmC sequencing; however, this limits our ability to measure both 5mC or 5hmC from the same genomic loci^19^. Another method, SIMPLE-seq, employs reactions to convert and detect 5mC and 5hmC in the genome; however, the efficiencies of these conversion reactions are less than 90%, resulting in a fraction of 5mC and 5hmC sites being incorrectly detected as unmethylated cytosines and reducing the accuracy of estimating 5mC/5hmC levels in the genome^20^. Similarly, DARESOME is another recent single-cell method that detects 5mC and 5hmC in addition to unmodified cytosines, using a series of restriction enzyme reactions to ligate different barcoded adapters to unmodified cytosine, 5mC, and 5hmC; however, a limitation of the method is that all three adapters have similar 3’ overhangs, potentially leading to incorrectly ligated adapters and increased false positive detection^21^. Furthermore, these methods are also unable to integrate these epigenetic measurements with quantification of the transcriptome from the same cell, thereby limiting our understanding of how turnover of the methylome directly impacts gene expression and cell state transitions. Therefore, in this study, we developed scMHT-seq, which employs two orthogonal restriction enzymes to profile 5mC and 5hmC with high specificity and combines it with molecular barcoding to also capture the transcriptome of the same cell in a one-pot assay to directly link DNA methylation dynamics to cell identity. We apply scMHT-seq to interrogate germ cell development at different time points to identify the modes of demethylation responsible for the erasure of the methylome *in vitro* and the relationship between the global methylation status and cell state. Finally, we demonstrate that integrated methylome and transcriptome sequencing in scMHT-seq enables us to identify potential factors involved in the genome-wide erasure of DNA methylation and maturation of germ cells, and suggests strategies to accomplish *in vitro* gametogenesis in the future.

## RESULTS

### scMHT-seq can jointly profile 5mC, 5hmC, and the transcriptome in single cells

To jointly profile both 5mC and 5hmC in single cells, we used two restriction enzymes that specifically recognize these two cytosine modifications^22–24^. We have previously developed single-cell methods using these restriction enzymes to strand-specifically quantify either 5hmC or 5mC in single cells^8,13,15,18,25^. To simultaneously profile 5mC, 5hmC, and mRNA from the same cells in scMHT-seq, we developed the following strategy: single cells are sorted into 384-well plates and the mRNA is reverse transcribed using polyT primers containing a cell-/RNA-specific barcode, unique molecule identifier (UMI), 5’ Illumina adapter and T7 promoter, followed by second strand synthesis to generate cDNA. Next, to identify 5hmC sites, genomic DNA is first treated with β-glucosyltransferase to glucosylate 5hmC marks, and subsequently, the restriction enzyme AbaSI is added which recognizes glucosylated 5hmC and generates double-stranded breaks with a 2 nucleotide 3’ overhang 11-13 nucleotides downstream of 5hmC^13,24,25^. Following this step, genomic DNA is treated with protease and the restriction enzyme MspJI is added, which recognizes 5mC sites in the genome and generates double-stranded breaks with a 4 nucleotide 5’ overhang 16 nucleotides downstream from 5mC^15,18,23^. Due to the orthogonal cutting modality of the two enzymes – with AbaSI generating 3’ overhangs and MspJI generating 5’ overhangs – two distinct sets of barcoded adapters are specifically ligated to AbaSI- and MspJI-generated cut sites. These two adapter sets contain a T7 promoter, 5’ Illumina adapter, and a 5mC- or 5hmC-specific cell barcode. As the cDNA and ligated gDNA molecules are tagged by cell- and molecule-of-origin-specific barcodes, they are pooled and amplified by *in vitro* transcription (IVT), followed by Illumina library preparation, as described previously, to enable joint profiling of 5mC, 5hmC and mRNA from single cells (Fig. 1b)^8,13,15,18,25–27^. Finally, as the two restriction enzymes AbaSI and MspJI generate sticky-end cut sites, and as IVT results in the directional amplification of molecules, the sequencing data can be used to computationally infer the strand-specificity of the two epigenetic modifications, as described previously^13,15,18^.

To benchmark the method, we first applied scMHT-seq to human embryonic stem cells (hESCs) and found that the efficiency of detecting unique 5mC and 5hmC sites in scMHT-seq was similar to those detected individually in scMspJI-seq and scAba-seq, and the number of unique mRNA transcripts detected by scMHT-seq was similar to other single-cell methods (Fig. 1c-d and Supplementary Fig. 1-3)^13,15,26^. As the detection of 5mC and 5hmC relies on restriction enzymes, we next wanted to ensure that we have high specificity and accuracy in detecting these two epigenetic modifications in scMHT-seq. Therefore, we evaluated the extent to which reads could potentially be assigned incorrectly to the other mark. In scAba-seq, we found that 3.23% of reads had a cytosine 16 nucleotides downstream from the cut site and could potentially be derived from MspJI cutting a methylated cytosine at that position (Supplementary Fig. 4a). However, as MspJI is absent in the scAba-seq workflow, this level of reads correspond to background noise arising from the random occurrence of cytosines at a specific position in the genome. Consistent with these observations, we found that 3.19% of reads in the 5hmC datasets in scMHT-seq could potentially arise from a 5hmC-specific adapter capturing an MspJI cut site; however, this is similar to the background noise established in the scAba-seq control dataset, suggesting minimal 5hmC miscalls (Supplementary Fig. 4a). Similarly, 5.19% and 4.02% of reads in 5mC datasets in scMspJI-seq and scMHT-seq, respectively, could potentially arise from 5hmC sites (Supplementary Fig. 4b). Overall, these results show that the false positive rates in scMHT-seq are low and similar to control datasets in scAba-seq and scMspJI-seq, demonstrating that there is minimal crosstalk between 5mC and 5hmC measurements in scMHT-seq. We next benchmarked the genome-wide distribution of 5mC and 5hmC obtained from scMHT-seq. For example, we observed the expected hypomethylation around CpG islands, consistent with 5mC measurements from scMspJI-seq and 5hmC measurements from scAba-seq (Supplementary Fig. 5a-b)^1^. Further, compared to scAba-seq and scMspJI-seq, we observed that scMHT-seq displayed a similar distribution of 5mC and 5hmC at gene promoters and gene bodies (Supplementary Fig. 5c-d). More generally, we observed that while scMHT-seq slightly undersamples 5mC and 5hmC sites within high density CpG regions of the genome, possibly due to over-digestion by the two restriction enzymes at these dense CpG regions, overall, we find that the distribution of CpG sites covered by scMHT-seq is similar to that found in the genome and that detected by scMspJI-seq and scAba-seq (Supplementary Fig. 6).

We next explored the relative distribution of 5mC and 5hmC in individual cells. Compared to 5mC, 5hmC is known to occur at much lower frequencies in most cell types, and we found that, on average, 5hmC levels in single cells are 3.1% of 5mC in the CpG context, consistent with previous estimates in hESCs (Fig. 2a)^28^. Further, we investigated the distribution of these epigenetic marks on individual DNA strands in single cells. As scMHT-seq is a strand-specific method, we assessed the strand asymmetry in the distribution of 5hmC and 5mCpG in single cells using a metric that we have previously used, called strand bias, defined over a genomic region as the number of epigenetic marks on the plus strand divided by the total detected on both strands. Thus, a strand bias of 0.5 indicates that both DNA strands have equal levels of the epigenetic mark whereas deviations from 0.5 indicate differences in the levels of the epigenetic mark between the older inherited DNA strand and the most recently synthesized new DNA strand. We found that the 5hmC strand bias was broadly distributed between 0.1 and 0.9 while the 5mCpG strand bias showed a tight distribution centered around 0.5 (Fig. 2b). This distribution is similar to our previous observations of 5hmC and 5mC strand bias in mouse embryonic stem cells (Supplementary Fig. 7). Furthermore, these results are consistent with the known mechanism of DNMT1-mediated inheritance of 5mCpG from old to new DNA strands, and the lack of inheritance of 5hmC and slow kinetics of TET activity that results in new DNA strands having lower levels of 5hmC compared to old DNA strands. Finally, the strand bias distributions of 5hmC and 5mCpG obtained from scMHT-seq are in close agreement to those obtained individually from either scAba-seq or scMspJI-seq, further validating that scMHT-seq can distinguish between 5hmC and 5mC with high accuracy and specificity (Fig. 2c and Supplementary Fig. 8).

**Figure 2.**
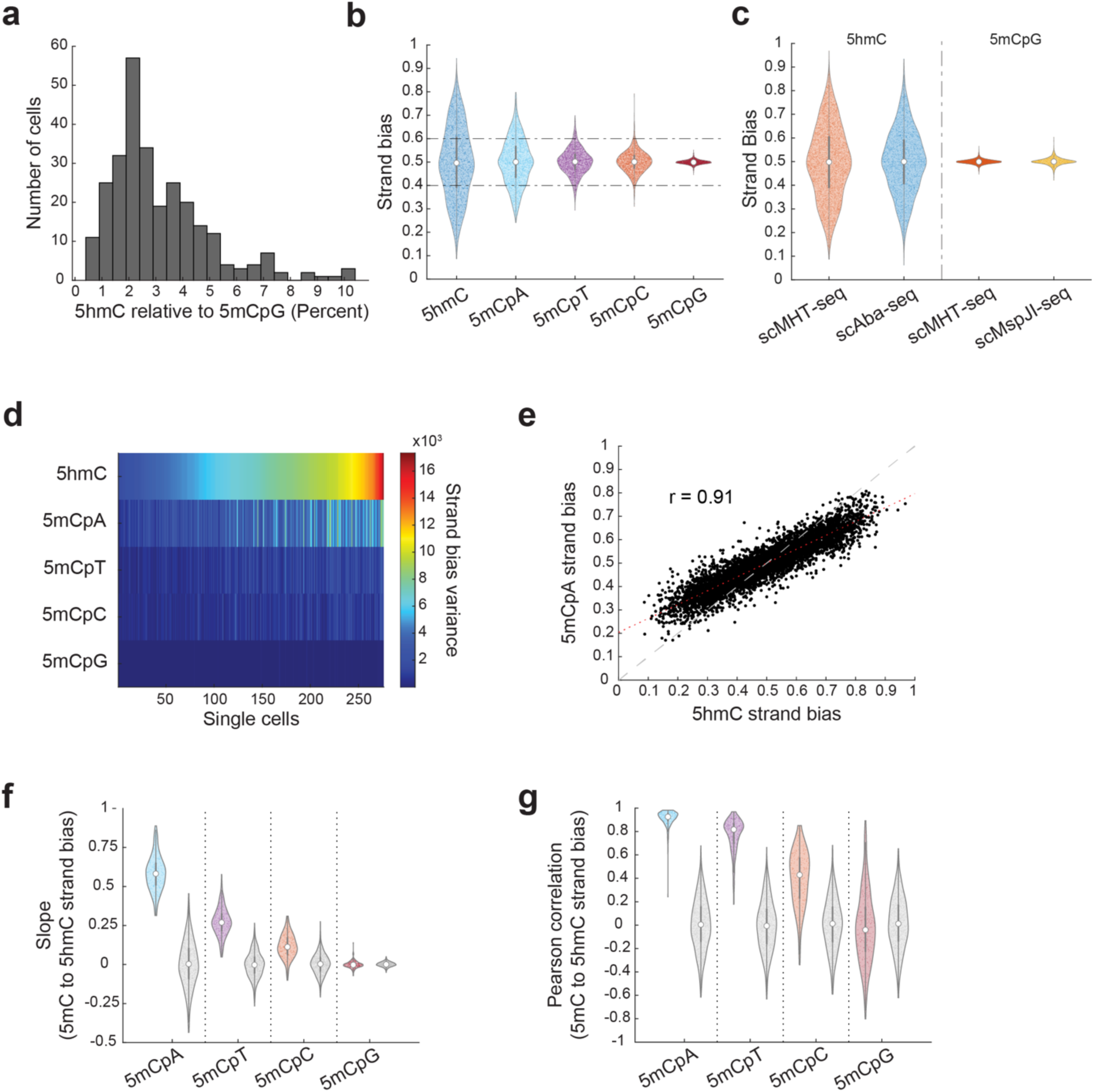
Dynamics between 5hmC, 5mCpG, and non-CpG methylation in individual cells. (a) Distribution of the ratio of 5hmC to 5mCpG sites detected by scMHT-seq in individual cells. (b) Violin plots of chromosome-wide strand bias distribution for 5mCpG, non-CpG methylation, and 5hmC. Each point represents an individual chromosome in single cells. (**c**) Chromosome-wide strand bias distributions for 5hmC and 5mCpG, measured with scAba-seq, scMspJI-seq, and scMHT-seq. Each point represents an individual chromosome in single cells. Similar strand bias distributions obtained from scMHT-seq and the individual methods indicate minimal crosstalk between detecting 5mC and 5hmC in scMHT-seq. (**d**) Heatmap of the variance in the strand bias distribution from all autosomal chromosomes of a single cell for 5mCpG, non-CpG methylation, and 5hmCpG. Individual cells are ordered based on those that display the lowest to the highest strand bias variance in 5hmC. (**e**) Comparison of 5mCpA strand bias to 5hmC strand bias for individual chromosomes across all single H9 cells profiled by scMHT-seq. The best fit line is shown in dotted red. (**f,g**) Comparison of 5hmC strand bias to 5mCpG and 5mCpH strand bias for the same chromosome in individual cells. Each point indicates the slope (f) or Pearson correlation (g) of a single cell. Gray dots within gray violin plots indicate an *in silico* cell, where the strand bias of each DNA modification on a chromosome was randomly sampled from the experimental dataset.

After establishing and benchmarking scMHT-seq, we next wanted to use the method to quantify the dynamics of methylation and hydroxymethylation in single cells. Specifically, we hypothesized that strand-specific quantification of non-CpG methylation could potentially be used infer the dynamics of *de novo* methylation, analogous to our previous work where we showed that strand-specific measurements of 5hmC could be used to gain insights into TET dynamics^13,25^, particularly as the DNA methylation maintenance machinery is known to have a strong preference for hemi-methylated CpG dinucleotides only^29^. Therefore, similar to 5hmC, newly synthesized DNA strands initially lack non-CpG methylation, and if the net rates of accumulation of non-CpG methylation are slow relative to the cell cycle, the old inherited DNA strand for any chromosome will contain higher levels of non-CpG methylation relative to the new DNA strand. In support of this mechanism, and consistent with our previous observations for 5hmC, we discovered high levels of strand-specific asymmetry in the levels of non-CpG methylation, resulting in broad strand bias distributions for non-CpG methylation (Fig. 2b). In agreement with these observations, we also found that the variance in strand bias between chromosomes of individual cells was higher for 5hmC and non-CpG methylation compared to 5mCpG (Fig. 2d). Furthermore, across individual chromosomes, we interestingly observed that 5hmC strand bias and non-CpG methylation strand bias were highly correlated (Pearson’s correlation, *r* = 0.91, 0.78, and 0.42 for 5mCpA, 5mCpT, and 5mCpC, respectively) (Fig. 2e and Supplementary Fig. 9). This coupling of 5hmC and non-CpG methylation strand bias arises because these epigenetic marks are inherited together on the old chromosome strand, thereby resulting in highly correlated transmission of 5hmC and non-CpG methylation. In contrast, the 5hmC strand bias was found to be uncorrelated with 5mCpG strand bias (Pearson’s correlation, *r* = 0.05), consistent with the known role of DNMT1 in copying 5mCpG from old to new DNA strands, thereby making the genome-wide distribution of 5mCpG largely independent of the age of the DNA strand. In summary, joint profiling of 5hmC and 5mC using scMHT-seq shows that unlike 5mCpG, the genome-wide strand-specific distributions of 5hmC and non-CpG methylation in single cells are closely linked and directly related to the age of the old inherited DNA strands.

Our comparison of non-CpG methylation to 5hmC strand bias interestingly also revealed that the slope of the best fit line was less than 1. For example we found that on average, the slope of 5mCpA *vs.* 5hmC strand bias was 0.59, implying that the strand bias of 5mCpA is always closer to 0.5 than that of 5hmC, and suggesting that the net rate of accumulation of 5mCpA on new DNA strands compared to old DNA strands due to the action of *de novo* methyltransferases and TET proteins is faster than that of 5hmC due to the action of TET proteins (Fig. 2e). To further investigate these observations, we estimated these correlations and slopes for individual cells, and compared the results to *in silico* derived cells where the chromosome-wide distribution of these epigenetic marks were randomly sampled from a background distribution derived from the experimental data (Fig. 2f-g). While the randomly generated *in silico* cells lacked correlation between 5hmC and 5mCpA strand bias on average, our experimental data showed strong Pearson’s correlations and slopes in all individual cells (mean *r* = 0.91 and mean slope = 0.58) (Fig. 2f-g), further demonstrating that 5hmC accumulates at a slower rate than 5mCpA in individual hESCs. To further validate these results, we performed stochastic modeling to estimate the turnover rates for the different epigenetic marks, using a model we previously developed^13^. In agreement with our results, we found the turnover rate of 5mCpA to be higher than 5hmC (Supplementary Fig. 10). Overall, these results show that scMHT-seq can accurately profile 5hmC, 5mCpG, and non-CpG methylation simultaneously from single cells with results that closely match individual measurements in single cells. Finally, these results show that strand-specific quantification of 5mC and 5hmC from the same cell in scMHT-seq can be used to estimate the kinetics of accumulation and loss of the methylome in mammalian cells.

### scMHT-seq identifies DND1 and SOX15 as potential factors involved in the global passive erasure of DNA methylation during hPGCLC maturation

After establishing scMHT-seq and validating that it can quantitatively profile DNA methylation, DNA hydroxymethylation and transcriptomes from single cells, we next wanted to apply the method to gain deeper insights into dynamic methylome reprogramming in the human germline.

Preimplantation development and the maturation of primordial germ cells (PGC) during early mammalian embryogenesis is characterized by dramatic global erasure of the methylome that is driven both by active demethylation, arising from the conversion of 5mCpG to 5hmCpG, as well as passive demethylation, arising from a reduction or lack of DNMT1-mediated 5mCpG maintenance activity (Fig. 1a)^1^. While this genome-wide erasure of DNA methylation in hPGCs is critical for the development of the germline *in vivo*, recent efforts to mimic this process *in vitro* by generating human PGC-like cells (hPGCLCs), with the long-term goal of using these technologies to accomplish *in vitro* gametogenesis, has been limited to cells with features akin to recently specified PGCs which have not undergone complete global DNA demethylation^10^. Recently, we developed an *in vitro* system that supports longer term culture of hPGCLCs where we observed partial demethylation (Fig. 3a)^11^. However, the contribution of active and passive demethylation to the global loss of the methylome during hPGCLC maturation remains unclear and the effect of 5mCpG erasure on cell identity is currently unknown. As this system is potentially composed of a heterogeneous population, bulk techniques can only provide limited insights; therefore, we hypothesized that quantifying 5mC, 5hmC and mRNA from the same cell using scMHT-seq will allow us to map the dynamics of the erasure of the methylome and enable us to directly link epigenetic reprogramming to cell identity during hPGCLC development. Thus, we applied scMHT-seq to hPGCLCs 4 days after induction (denoted D4) and hPGCLCs cultured for an additional 10 or 21 days in long-term culture conditions (denoted D4C10 and D4C21, respectively) (Fig. 3a).

**Figure 3.**
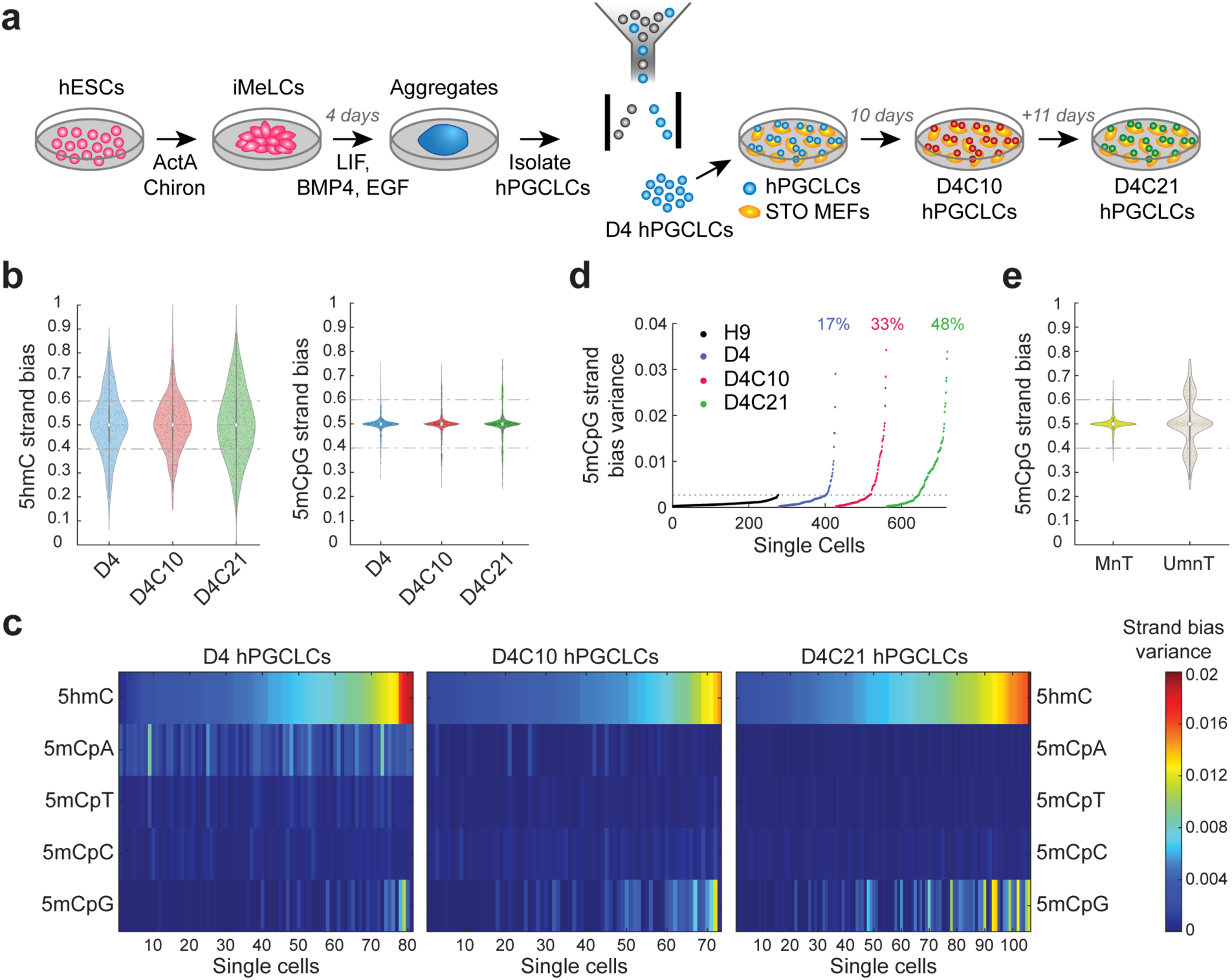
Long-term culture of hPGCLCs results in the emergence of globally heterogenous and partially demethylated hPGCLCs. (**a**) Schematic for generating hPGCLCs in extended culture. hESCs are first differentiated to iMeLCs and four days after induction, hPGCLCs are isolated from aggregates by FACS and cultured for an additional 10 days or 21 days under extended culture conditions. (**b**) 5hmC (left) and 5mCpG (right) strand bias of hPGCLCs at different stages of culture. Data points represent individual chromosomes in single cells. (**c**) Heatmaps of the variance in the strand bias distribution from all autosomal chromosomes of a single cell for 5mCpG, non-CpG methylation and 5hmCpG, separated by hPGCLC culture conditions. Individual cells are ordered based on those that display the lowest to the highest strand bias variance in 5hmCpG. (**d**) Comparison of the variance in the 5mCpG strand bias distribution of individual hPGCLCs derived from different time points in the extended culture. Single cell data were acquired with scMHT-seq or scMspJI-seq. Horizontal gray line indicates the maximum variance in 5mCpG strand bias observed in individual H9 hESCs, cells that are known to display DNMT1-mediated maintenance of DNA methylation. Cells above this line were those that were categorized as showing impaired DNA methylation maintenance (unmaintained or ‘UmnT’). Cells below this line were those that were categorized as displaying high DNA methylation maintenance (maintained or ‘MnT’). Percent values indicate percentage of hPGCLCs in the UmnT group for a given condition. (**e**) 5mCpG strand bias of ‘MnT’ and ‘Umnt’ hPGCLCs over all stages of culture. Data points represent individual chromosomes in single cells.

To assess the state of the methylome of hPGCLCs at all three time points, we first quantified strand bias in single cells. As with hESCs, we observed a wide unimodal 5hmC strand bias distribution and a high degree of cell-to-cell heterogeneity in 5hmC strand bias (Fig. 3b,c). However, unlike undifferentiated hESCs, we found that the 5mCpG strand bias distribution was wider with longer tails in all hPGCLC culture conditions (Fig. 3b). Notably, we discovered that a subset of cells exhibited high variance in 5mCpG strand bias which coincided with high variance in 5hmC strand bias for the same cells, suggesting that a subset of hPGCLCs in all conditions have reduced 5mCpG on new DNA strands compared to old strands, implying that these cells are undergoing genome-wide passive demethylation due to impaired DNMT1-mediated maintenance methylation activity (Fig. 3c). Furthermore, in support of these results, we found that 5hmC and 5mCpG strand bias across chromosomes in individual cells were strongly correlated at all three time points, clearly highlighting the existence of passive demethylation in this system, consistent with our previous observations (Supplementary Fig. 11a)^11^. Interestingly, compared to hESCs, we generally also observed a significantly weaker correlation and slope between 5hmC and non-CpG methylation, indicating distinct non-CpG methylation dynamics in hPGCLCs (Fig. 1f-g, Supplementary Fig. 11a,b). Next, to identify the subset of hPGCLCs that are passively demethylating, we compared D4, D4C10 and D4C21 hPGCLCs to hESCs and classified cells as those that display high (maintained or ‘MnT’) or impaired (unmaintained or ‘UmnT’) DNA methylation maintenance. Strikingly, we find that a substantial fraction of hPGCLCs are passively demethylating at all three time points, with a larger proportion of hPGCLCs passively demethylating at longer culturing times (Fig. 3d,e). Overall, these results from scMHT-seq show that hPGCLCs in this *in vitro* culture system show dramatic cell-to-cell heterogeneity with two population of cells displaying distinct global methylome profiles (Fig. 3e).

We then focused on the transcriptome of these cells to identify three distinct transcriptional clusters. We found that D4 hPGCLCs cluster separately from those in extended culture (denoted transcriptional cluster ‘D4’), and that within the long-term culture conditions D4C10 and D4C21, two distinct transcriptional states can be observed (denoted transcriptional clusters ‘LT1’ and ‘LT2’) (Fig. 4a). Interestingly, we found that the transcriptional cluster assignment of LT1 and LT2 appeared to be independent of whether hPGCLCs were cultured for 10 or 21 days in long-term conditions (Fig. 4a). While expression of key hPGC genes, such as SOX17, PRDM1 (also known as BLIMP1), TFAP2C and NANOS3, were found in hPGCLCs from all three time points, there were key differences between the three transcriptional clusters, with the largest changes observed between the D4 and long-term culture groups LT1 and LT2, and these transcriptional programs agreed well with prior bulk RNA-seq and immunofluorescence results (Fig. 4b)^11^. Further, promoter and gene body 5mCpG levels decreased during the progression of hPGCLCs from D4 to LT1 and LT2, further confirming the ability of long-term culture conditions to maintain a germ cell state as they initiate genome-wide erasure of DNA methylation (Fig. 4c).

**Figure 4.**
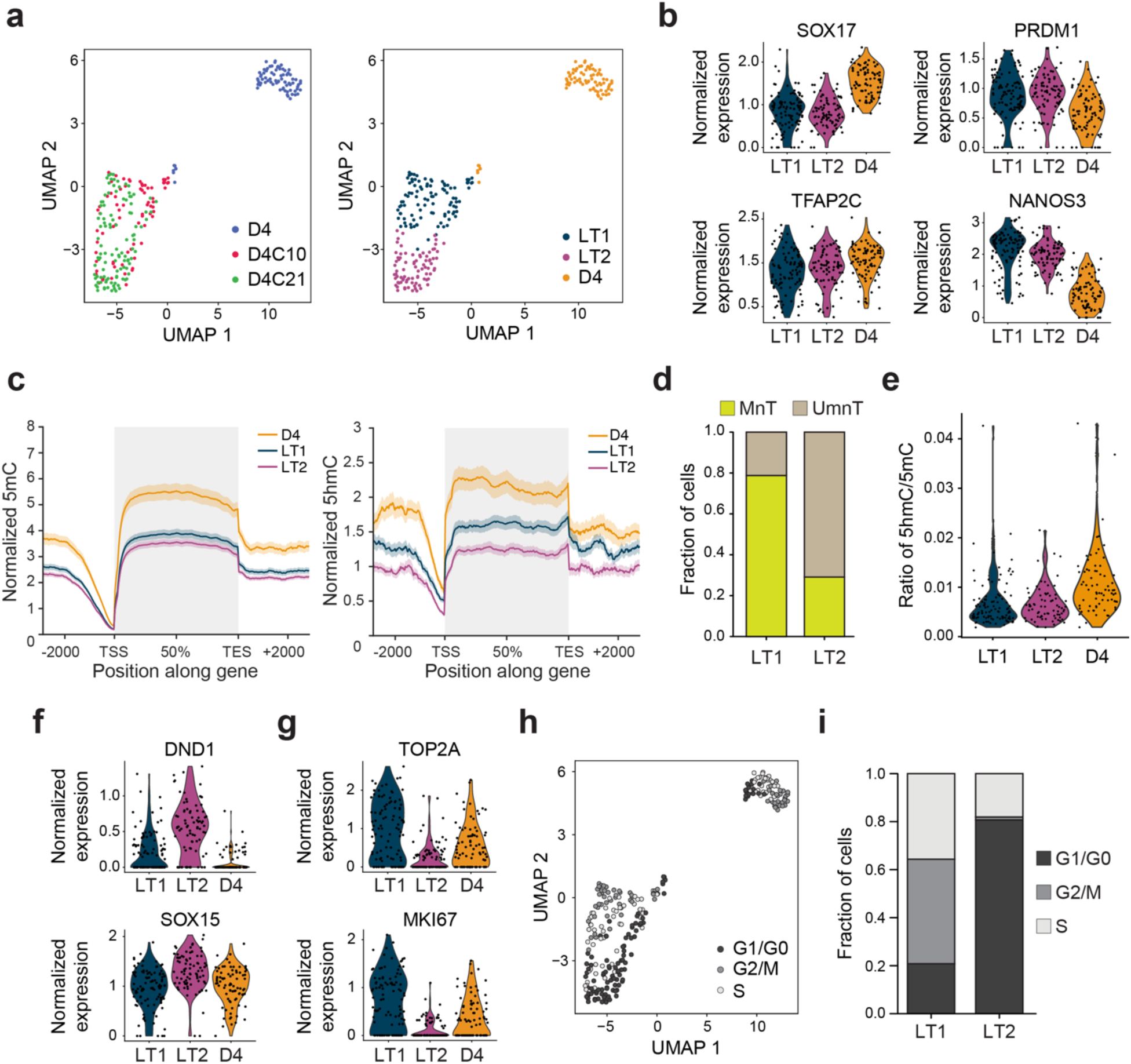
hPGCLCs in extended culture display two distinct transcriptional states that are directly associated with the genome-wide loss of DNA methylation. (**a**) UMAP projection of the transcriptome of hPGCLCs, grouped by culture conditions (*left*) or gene expression-based clustering (*right*). (**b**) Normalized expression of key hPGC genes SOX17, PRDM1 (also known as BLIMP1), TFAP2C, and NANOS3 in hPGCLCs derived from long-term cultures in D4, LT1, and LT2 transcriptional clusters. (**c**) Normalized 5mCpG (*left*) and 5hmC (*right*) profiles over promoters and gene bodies of hPGCLCs in D4, LT1, and LT2 transcriptional clusters. Counts of 5mCpG and 5hmC were normalized against their respective 5mCpG and 5hmC spike-in counts. Solid lines indicate mean normalized counts per genomic bin. Shaded regions indicate standard error of the mean, computed over 10,000 bootstrap samples. (**d**) Bar plots show the fraction of hPGCLCs in LT1 and LT2 transcriptional groups that display passive demethylation (unmaintained or ‘UmnT’) or high DNA methylation maintenance (maintained or ‘MnT’). (**e**) Ratio of 5hmC to 5mCpG marks detected in individual hPGCLCs from D4, LT1, and LT2 transcriptional groups. (**f,g**) Normalized expression of genes found to be differentially expressed between LT1 and LT2, with genes such as DND1 and SOX15 found to be critical for germ cell development and maturation (f), and genes such as TOP2A and MKI67 related to cell cycle progression (g). (**h**) UMAP projection of the transcriptome of hPGCLCs grouped by predicted cell cycle stage (G1/G0, G2/M, or S phase) (**i**) Predicted cell cycle phase of hPGCLCs in LT1 and LT2 transcriptional clusters.

While hPGCLCs in these long-term cultures maintained germline identity, we observed pronounced heterogeneity spanning both their epigenetic (MnT *vs.* UmnT) and transcriptional (LT1 *vs.* LT2) states. We hypothesized that these epigenetic and transcriptional states may be linked and mimic hPGC maturation *in vivo*. Strikingly, we discovered that the LT2 transcriptional state is highly enriched for UmnT cells, demonstrating that this particular gene expression program is associated with passively demethylating hPGCLCs (Fig. 4d). Furthermore, hPGCs *in vivo* initiate genome-wide erasure of DNA methylation after initial specification, suggesting that cells in the LT2 transcriptional group are potentially more mature hPGCLCs than those in LT1^3,30,31^.

After establishing that differential DNMT1-mediated maintenance methylation fidelity is associated with the two long-term culture transcriptional groups LT1 and LT2, we next focused on understanding if these hPGCLCs also undergo active demethylation, as germ cells *in vivo* have been shown to globally erase their methylome through a combination of passive and active demethylation^2–4,30–33^. Analysis of global 5hmC levels in individual cells showed similar distributions for both LT1 and LT2, and when compared to D4 hPGCLCs, the levels of 5hmC relative to 5mC in LT1 and LT2 hPGCLCs was low, suggesting that these long-term germ cells do not undergo active demethylation (Fig. 4e). In agreement with these results, we also found that LT1 and LT2 hPGCLCs had lower levels of 5hmC over promoters and gene bodies compared to D4 hPGCLCs (Fig. 4c). Finally, consistent with these observations, we found that 5hmC levels are similar in the Mnt and UmnT groups as well, indicating a lack of active demethylation (Supplementary Fig. 12). Taken together, these findings suggest that the hPGCLCs cultured in this system are not fully demethylated and primarily exhibit passive but not active demethylation. Finally, as the LT2 population potentially reflects a more advanced hPGCLC state compared to LT1, we wanted to identify the transcriptional signatures that are associated with this maturation process. The genes that are differentially expressed between the two transcriptional states within long-term culture, LT1 and LT2, are expressed at similar levels in LT1 and D4 hPGCLCs, suggesting that the LT1 population is transcriptionally closer to the D4 hPGCLCs, which have not experienced extended culture, compared to LT2 (Supplementary Fig. 13). Next, inspection of differentially expressed genes identified putative genes involved in PGC maturation and the initiation of global erasure of DNA methylation (Supplementary Fig. 13). Notably, we discovered that DND1 and SOX15 are expressed at higher levels in LT2 compared to LT1, two factors that have been shown to be important for germ cell development (Fig. 4f)^34–39^. For example, DND1 has been shown to be associated with mouse germ cell development, with a dramatic reduction in PGC numbers occurring after specification in mutants lacking wild-type protein expression^34^. Similarly, DND1 has also been implicated in human germ cell development, as DND1 knockout cells have a reduced capacity to differentiate into hPGCLCs^36^. Further, while SOX15 has been shown to be dispensable for initial human primordial germ cell establishment, it is potentially important for germ cell maintenance, as SOX15 depletion leads to reduced hPGCLCs in *in vitro* models^35^. In addition, DND1 expression in Xenopus directly regulates NANOS1, a key regulator of PGC fate in this organism^37,38^. Interestingly, DND1 has also been strongly implicated in the downregulation of active cell cycle genes, with DND1 mutants causing gonadal teratoma formation in mice^40^. Consistent with this role of DND1 in other species, we found that active cell cycle genes, such as TOP2A and MKI67, are downregulated in the LT2 hPGCLC population, which expresses DND1 at higher levels (Fig. 4f-g). In agreement with this observation, cell cycle analysis revealed that a larger fraction of cells in LT2 are non-cycling, likely at least in part due to high levels of DND1 (Fig. 4h-i). The observation that many LT2 cells are non-dividing while simultaneously displaying passive global demethylation suggests that these cells were dividing previously with impaired fidelity of DNMT1-mediated maintenance methylation (Fig. 4d,i). Overall, these results from scMHT-seq suggests that factors identified in this work, such as DND1 and SOX15, are potentially key regulators of the global erasure of DNA methylation and maturation of human germ cells, thereby providing strategies to advance hPGCLC development *in vitro* through targeted control of the expression of these genes (Supplementary Fig. 13).

## DISCUSSION

In this work, we have developed scMHT-seq, the first single-cell method to simultaneously profile both DNA methylation and DNA hydroxymethylation, together with the transcriptome from the same cell, in a single-tube assay that does not require physical separation of the nucleic acids prior to amplification, thereby minimizing losses and achieving efficiencies similar to individual measurements of scMspJI-seq, scAba-seq and scRNA-seq in single cells. Further, we show that the false positive rates of detecting 5mC and 5hmC in scMHT-seq is low and similar to those estimated in scMspJI-seq and scAba-seq. Importantly, scMHT-seq also enables genome-wide strand-specific measurements of 5mC and 5hmC, allowing quantification of methylation on sense and antisense strands, and providing additional insights into the mechanism driving DNA methylation erasure by deconvolving passive from active demethylation. Finally, we demonstrate that the strand-specific measurements of scMHT-seq in hESCs provides quantitative estimates of the kinetics of turnover of 5hmC, 5mCpG and non-CpG methylation on newly synthesized DNA strands during each cell cycle relative to older DNA strands.

Previously, the study of 5mC and 5hmC dynamics was primarily limited to computational integration of separate experimental datasets; however, such approaches are not able to directly interrogate the underlying rates of methylation and demethylation that regulate the turnover of the methylome in individual cells^7,41^. Therefore, more recently a few techniques have emerged that enable simultaneous profiling of 5mC and 5hmC. Our group (Dyad-seq) and others (Six-letter-seq) have developed techniques to profile 5mC and 5hmC from the same sample; however, these methods have not been scaled down to single cells^17,18^. In contrast, while Joint-snhmC-seq enables single-cell measurements, this technique requires splitting the genomic material into two halves; therefore, a key limitation of this method is that it cannot detect 5mC and 5hmC from the same genomic locus, thereby resulting in low-resolution coverage of the genome and an inability to quantify locus-specific relationships between 5mC and 5hmC^19^. Another recent method, SIMPLE-seq, detects 5mC and 5hmC in single cells by performing reactions to convert the methylated and hydroxymethylated cytosines; however, these reactions have a conversion rate of less than 90%, unlike bisulfite-based methods that have a conversion rate of unmethylated cytosines to uracils of greater than 98%, resulting in a significant fraction of 5mC/5hmC sites being incorrectly called as unmethylated cytosines, thereby offering less reliable estimates of the methylation and hydroxymethylation levels on a locus-specific or genome-wide scale^20^. To overcome these limitations, scMHT-seq offers an approach free of nucleobase conversion, and instead leverages the highly specific recognition activity of restriction enzymes to discriminate between 5mC and 5hmC in a single-tube reaction that does not require splitting the genomic DNA prior to amplification. DARESOME, another recent single-cell method uses restriction enzymes to simultaneously profile 5mC, 5hmC, and unmodified cytosines; however, compared to scMHT-seq which uses AbaSI and MspJI to recognize modified cytosines in CN20-23G and CNNR contexts, respectively, the DARESOME restriction enzymes HpaII and MspI recognize cytosines in a more limited sequence context of CCGG sites, and therefore sample considerably fewer genomic loci^21^. Importantly, unlike the other methods discussed above, scMHT-seq enables capturing the transcriptome, together with 5mC and 5hmC, from the same cell. Therefore, this multiomic single-cell sequencing strategy enables us to directly link methylation dynamics to gene expression and cell state in individual cells. Overall, scMHT-seq provides an accurate, sensitive and high-throughput approach to reliably capture 5mC, 5hmC and mRNA in single cells, thereby offering a comprehensive view of the cytosine methylation cycle in complex and heterogeneous cell populations.

Finally, we applied scMHT-seq to an hPGCLC extended culture system we had previously developed to map the global erasure of DNA methylation in this *in vitro* system^11^. *In vivo*, hPGCs undergo dramatic genome-wide loss of DNA methylation, a key event that plays a central role in the maturation of germ cells and the eventual differentiation towards gametes; however, mimicking this global epigenetic reprogramming event *in vitro* remains a challenge, thereby limiting our ability to accomplish gametogenesis in a dish. Therefore, we used scMHT-seq to quantify the extent and mechanism through which hPGCLCs demethylate in our *in vitro* system. In extended culture, we observed two transcriptionally distinct groups of hPGCLCs, and leveraged scMHT-seq to excitingly find that one of these groups primarily consists of passively demethylating cells that represent a more mature germ cell state. Importantly, combined measurements of the methylome and transcriptome led us to identify genes that are differentially expressed in the demethylating population, and thereby enabled us to find factors that may potentially be involved in the erasure of DNA methylation and germ cell maturation. In particular, we discovered that cells in a demethylated state expressed high levels of DND1 and SOX15, two genes that have previously been shown to be critical for PGC maturation in various species, suggesting a key role for these factors in human germ cell development as well. Recently, CRISPR interference (CRISPRi) screens have also been used to identify factors important for initial hPGCLC specification; however, without measurements of the methylome, it is unclear if these genes affect DNA methylation dynamics and hPGCLC maturation^42–44^. Overall, these results demonstrate that scMHT-seq not only enables detailed insights into the mechanisms driving DNA methylation dynamics but also through the joint measurement of 5mC, 5hmC and mRNA directly help identify the interplay between factors that regulate the methylome and the impact of the epigenome on transcription and cell states.

## MATERIALS AND METHODS

### Mammalian cell culture

Culturing human embryonic stem cells: All cells were maintained in cell culture incubators at 37°C and 5% CO2. H9 human embryonic stem cells were grown feeder-free on plates coated with Matrigel (Fisher Scientific, 08-774-552) in mTeSR1 media (STEMCELL Tech., 85850) as described previously^8^. For sorting individual cells into 384-well plates for use in scMHT-seq, a single cell suspension is first generated using 0.25% trypsin-EDTA, and the trypsin is subsequently inactivated using serum containing media. The cells are then washed with 1X PBS and passed through a cell strainer before being single-cell sorted using a FACS instrument into 384-well plates.

Generation of hPGCLCs and extended culture: UCLA2 human embryonic stem cells were cultured and induced into hPGCLCs via an incipient mesoderm-like cell intermediate and the generation of disorganized 3D aggregates, as previously described^45^. After 4 days in 3D culture, hPGCLCs were sorted and cultured in extended culture conditions containing FR10 medium, as described previously^11^. Following extended culture, TRA-1-85 positive single hPGCLCs were isolated into each reaction well of a 384-well plate for scMHT-seq, as described previously^11^.

### scMHT-seq

Four microliters of Vapor-Lock (QIAGEN, 9881611) was added to each well of a 384-well plate. Thereafter, all the following dispensing steps were performed using a Nanodrop II liquid-handling robot (BioNex Solutions). Next, 100 nL of uniquely barcoded reverse transcription primer (7.5 ng/uL) was added (primers described in Grun et al. with the exception that a 6 bp UMI was used) to each well^46^. Just prior to cell sorting, 100 nL of lysis buffer (0.175% IGEPAL CA-630, 1.75 mM dNTPs (NEB, N0447S), 1:1,250,000 ERCC RNA spike-in mix (Ambion, 4456740) and 0.19 U RNAse inhibitor (Clontech, 2313A)) was added to each well. Single cells were sorted into each well using FACS and stored at -80°C. To begin processing, plates were heated to 65°C for 3 min and returned to ice. Thereafter, 150 nL of reverse transcription mix was added (0.7 U RNAaseOUT (Invitrogen, 10777-019), 2.33x first-strand buffer, 23.33 mM DTT and 3.5 U Superscript II (Invitrogen, 18064-071), and the plates were heated to 42°C for 75 min, 4°C for 5 min and 70°C for 10 min. Next, 1.5 uL of second-strand synthesis mix was added (1.23x second-strand buffer (Invitrogen, 10812-014), 0.25 mM dNTPs (NEB, N0447S), 0.14 U *E. coli* DNA Ligase (Invitrogen, 18052-019), 0.56 U *E. coli* DNA Polymerase I (Invitrogen, 18010-025), and 0.03 U RNase H (Invitrogen, 18021-071)), and the plates were then incubated at 16°C for 2 hours.

Following this step, 650 nL of protease mix (6 µg protease (Qiagen, 19155) and 3.85x NEBuffer 4 (NEB, B7004S)) was added to each well, and the plates were heated to 50°C for 15 h, 75 °C for 20 min and 80°C for 5 min. 500 nL of 5mC glucosylation mix (1 U T4-BGT (NEB, M0357L), 6x UDP–glucose, 200 fg of mouse brain spike-in DNA (used for estimating sequencing efficiency) (VWR, 76020-078) and 1x NEBuffer 4) was then added to each well, and the plates were incubated at 37°C for 16 h. Next, 500 nL of protease mix (2 µg protease and 1× NEBuffer 4) was added to each well, and the plates were incubated at 50°C for 3 h, 75°C for 20 min and 80°C for 5 min. After the second protease step, 500 nL of glucosylated 5hmC digestion mix was added (1x NEBuffer 4 (NEB, B7004S), 1 U AbaSI (NEB, R0665S)) to each well and the plates were incubated at 25°C for 90 minutes, and 65°C for 25 minutes. Next, 250 nL of protease mix (2 µg protease (Qiagen, 19155), and 1x NEBuffer 4) was added to each well, and the plates were heated to 50°C for 3 hours, 75°C for 20 minutes, and 80°C for 5 minutes. Next, 500 nL of MspJI digestion mix (1x NEBuffer 4, 9.5x enzyme activator solution, and 0.1 U MspJI (NEB, R0661L)) was added to each well and the plates were incubated at 37°C for 4.5 hours, and 65°C for 25 minutes. To each well, 200 nL of uniquely barcoded 20 nM phosphorylated scAba-seq compatible double-stranded adapters were added^13^. Then to each well, 120 nL of uniquely barcoded 125 nM phosphorylated scMspJI-seq compatible double-stranded adapters were added^15^. The sequences of scAba-seq and scMspJI-seq compatible double-stranded adapters have previously been reported by us^13,15^. Next, 680 nL of ligation mix (1.47x T4 ligase reaction buffer, 6.99 mM ATP (NEB, P0756L), and 140 U T4 DNA ligase (NEB, M0202M)) was added to each well, and the plates were incubated at 16°C for 16 hours. After ligation, reaction wells receiving different barcodes were pooled using a multichannel pipette, and the oil phase was discarded. Library preparation and Illumina sequencing for mRNA enriched and non-mRNA enriched samples was performed as described previously^8^. Finally, the individual scMspJI-seq and scAba-seq libraries that were used as controls for comparison to scMHT-seq were prepared as described previously^13,15^.

### scMHT-seq analysis pipeline

All sequencing reads were trimmed to 76 bases. Then 5mC, 5hmC, and transcriptome-based reads were separated based on feature-specific barcodes using custom Perl scripts (see Supplementary File 1). Thereafter, the scMHT-seq data analysis was performed as described previously in Sen *et al.* and Mooijman *et al*.^13,15^. To map reads to the human and mouse genome, hg19 and mm10 were used, respectively, with 5mC marks attributed to the mouse genome representing spike-in detections. The transcriptome analysis pipeline was previously described by us in Chialastri *et al*^8^. Each feature – 5mC, 5hmC or mRNA – was separately analyzed for data quality. If at least 30,000 5mC, 300 5hmC, 4,000 unique transcripts and 1,000 genes were detected in a cell, it was considered successfully sequenced in all features. In some cases, a cell only contained high quality information for one or two of these features; in such instances the cell was used in the analysis only when quantifying that feature.

### Estimating false positive rates

The false positive rate per single cell was defined as the fraction of reads where the unintended mark is called either jointly with the intended mark or on its own. Separated 5mC- and 5hmC-specific data (that is, sequencing reads already assigned as 5mC or 5hmC based on feature-specific barcodes) was searched to count the number of reads containing putative 5mCpG (A), reads containing putative 5hmCpG (B), and reads containing putative sites for both 5mCpG and 5hmCpG (C). The false positive rate for scAba-seq was A/(A+B-C). The false positive rate for scMspJI-seq was B/(A+B-C).

### Strand bias analysis

5hmC and 5mC strand biases were calculated per chromosome or for a region within a chromosome for each single cell. Strand bias is defined as the number of 5hmC or 5mC marks on the sense strand divided by the total number of 5hmC or 5mC marks on both strands.

### Gene expression analysis

The standard analysis pipeline in Seurat (version 3.1.5) was used for single-cell RNA-seq normalization and analysis^47^. Cells where more than 1,000 genes and more than 4,000 unique transcripts, as well as less than 20% ERCC spike-ins were detected, were used for downstream analysis. The default NormalizeData function was used to log normalize the data. Thereafter, principal components were obtained from the 2,000 most variable genes and the elbow method was used to determine the optimal number of principal components used in clustering. UMAP based clustering was performed by running the following functions, FindNeighbors, FindClusters, and RunUMAP. After clustering, cell types were assigned to groups using known expression markers. To identify differentially expressed genes (DEGs), the FindAllMarkers or FindMarkers function was used. The Wilcoxon rank sum test was used to classify a gene as differentially expressed, requiring a natural log fold change of at least 0.3 and an adjusted p-value of less than 0.01. Cell cycle analysis was performed as described in the Seurat cell cycle vignette using cell cycle genes derived previously^48^.

### Modeling turnover rates of 5hmC and 5mCpH in individual cells

The turnover of each modification (5hmC, 5mCpA, 5mCpT, 5mCpC) was modeled stochastically using chromosome-wide stand-specific scMHT-seq data. This stochastic model has been previously described by us in Mooijman *et al*.^13^. For each DNA modification, 100 simulations were performed.

## ACKNOWLEDGEMENTS

We would like to thank Zsófia Szegletes and other members of the Dey lab for helpful discussions and feedback. We thank Jennifer Smith at the Biological Nanostructures Laboratory in the California NanoSystems Institute (CNSI), supported by UCSB and UC Office of the President, for assistance with Illumina sequencing. Computational work was supported by the Center for Scientific Computing at CNSI and Materials Research Laboratory (MRL) at UCSB: an NSF MRSEC (DMR-1720256) and NSF CNS-1725797. S.E.W. was supported by a University of California Presidents Postdoctoral Fellowship and a Young Investigator Award from the Iris Cantor-UCLA Women’s Health Education & Research Center (NCATS UCLA CTSI Grant # UL1TR001881). This work was supported by the NIH grant 2R01HD079546 to A.T.C., and NIH grants R01HD099517 and R01HG011013, and an NSF CAREER grant (Award # 2339849) to S.S.D.

## AUTHOR CONTRIBUTIONS

Conceptualization, A.C. and S.S.D.; Methodology, A.C. and S.S.D.; Investigation, A.C., K.A.H., J.J.G., F.H., S.E.W.; Formal Analysis, A.C. and K.A.H.; Writing – Original Draft, A.C.; Writing – Review & Editing, A.C., K.A.H, and S.S.D.; Funding Acquisition, A.T.C. and S.S.D.; Resources, A.T.C. and S.S.D.; Supervision, S.S.D.

## ETHICS DECLARATION

The use of human embryonic stem cells has been approved by the UC Irvine Human Stem Cell Research Oversight Committee (Protocol #105) and the UCLA Embryonic Stem Cell Research Oversight Committee (Protocol #2006-005-18B).

## DATA AVAILABILITY

Sequencing data has been deposited in the Gene Expression Omnibus (GEO) database accession code GEO: GSE280112.

## CODE AVAILABILITY

Custom codes for analyzing scMHT-seq are deposited on GitHub: (https://github.com/alexchialastri/scMHT-seq/).

