## Supplementary Information for "Joint profiling of 5mC, 5hmC, and the transcriptome in single cells identifies factors responsible for genome-wide DNA methylation erasure in human primordial germ cell maturation"

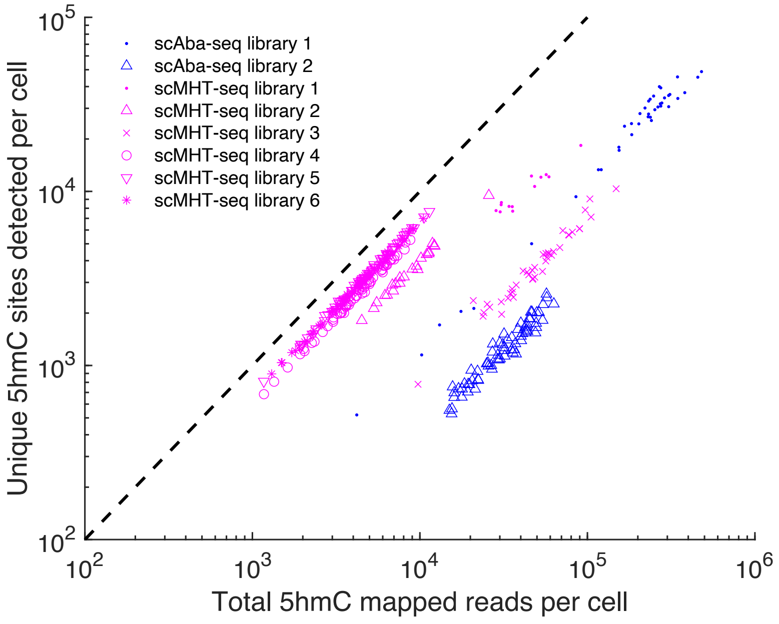


**Supplementary Figure 1 | Coverage of unique 5hmC sites detected per cell in scMHT-seq and scAba-seq.** The number of unique 5hmC sites detected per cell are shown as a function of the sequencing depth for scAba-seq and scMHT-seq. These experiments were performed on H9 hESCs. The number of unique 5hmC sites detected per cell increases with sequencing depth, suggesting that more unique sites could be detected per cell by sequencing the Illumina libraries deeper. Libraries closer to the dotted diagonal line indicate a higher complexity of unique 5hmC sites per sequenced read in the library. The complexities of the sequencing libraries are similar for both scAba-seq and scMHT-seq.

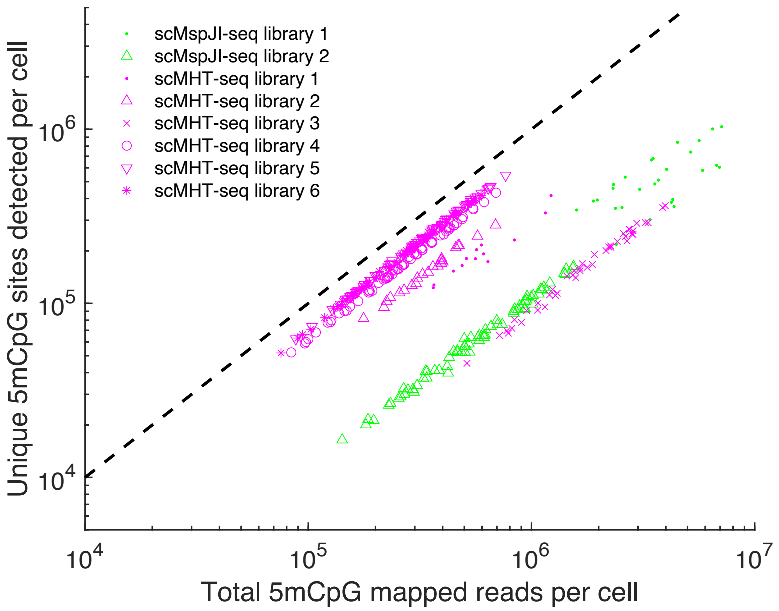


**Supplementary Figure 2 | Coverage of unique 5mCpG sites detected per cell in scMHT-seq and scMspJI-seq.** The number of unique 5mCpG sites detected per cell are shown as a function of the sequencing depth for scMspJI-seq and scMHT-seq. These experiments were performed on H9 hESCs. The number of unique 5mCpG sites detected per cell increases with sequencing depth, suggesting that more unique sites could be detected per cell by sequencing the Illumina libraries deeper. Libraries closer to the dotted diagonal line indicate a higher complexity of unique 5mCpG sites per sequenced read in the library. The complexities of the sequencing libraries are similar for both scMspJI-seq and scMHT-seq.

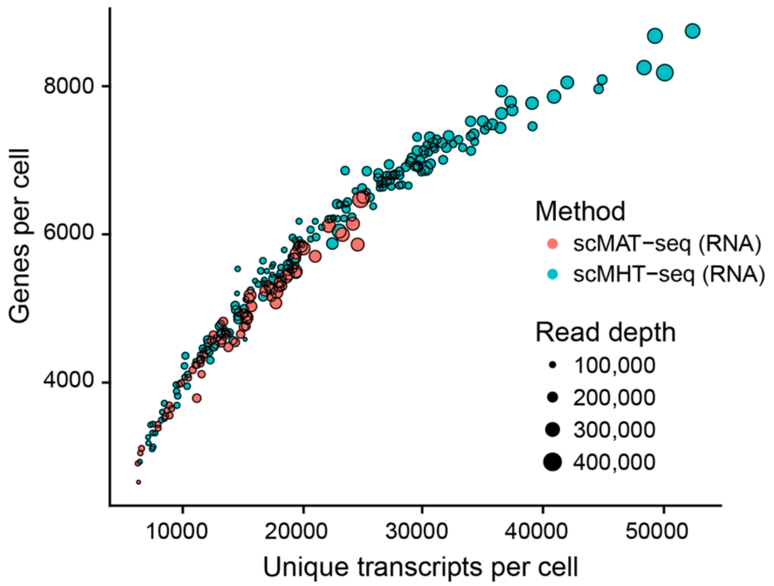


**Supplementary Figure 3 | Comparison of the single-cell transcriptomes obtained from scMHT-seq and scMAT-seq.** The single-cell transcriptomes of scMHT-seq were compared to those obtained from another single-cell multiomics method scMAT-seq. Both methods use T7-based linear amplification to detect unique transcripts in single cells. The number of unique genes detected per cell ranged from 2,648 to 6,505 for scMAT-seq and 2,923 to 8,748 for scMHT-seq. Similarly, the number of unique transcripts detected per cell ranged from 6,266 to 25,041 for scMAT-seq and 6,432 to 52,445 for scMHT-seq, suggesting that both methods have a similar efficiency in detecting unique mRNA molecules in individual cells. These experiments were performed on H9 hESCs.

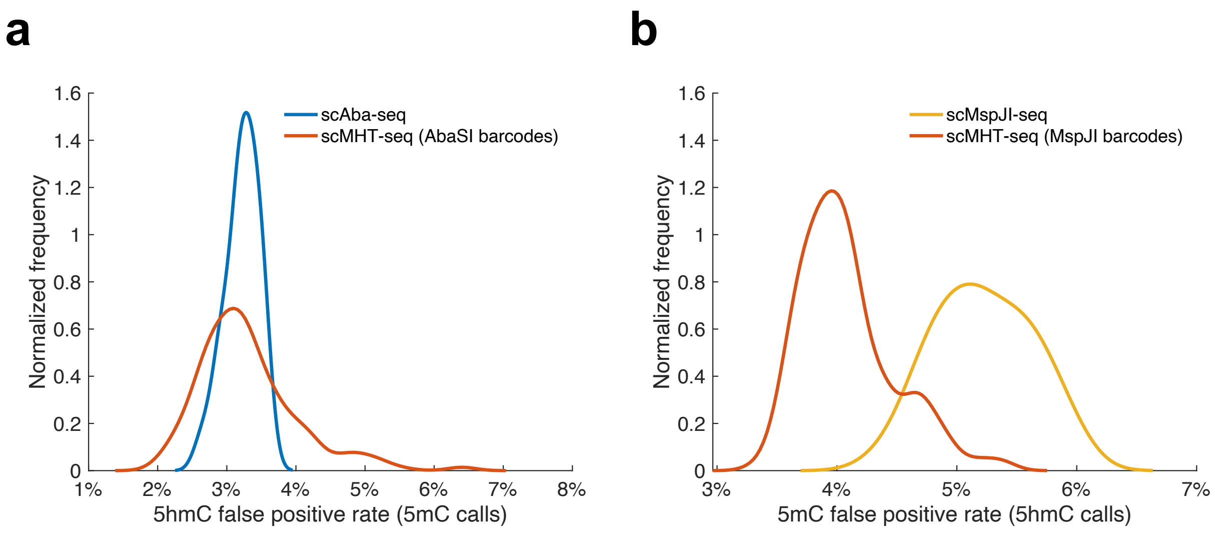


**Supplementary Figure 4 | Distribution of false positive detection rates of 5hmC and 5mC in scMHT-seq.** (**a**) Distributions of false positive detection rate of 5hmC per cell in scMHT-seq and scAba-seq, with medians of 3.19% and 3.23%, respectively. The distribution obtained from scAba-seq represents background noise. (**b**) Distributions of false positive detection rate of 5mC per cell in scMHT-seq and scMspJI-seq, with medians of 4.02% and 5.19%, respectively. The distribution obtained from scMspJI-seq represents background noise.

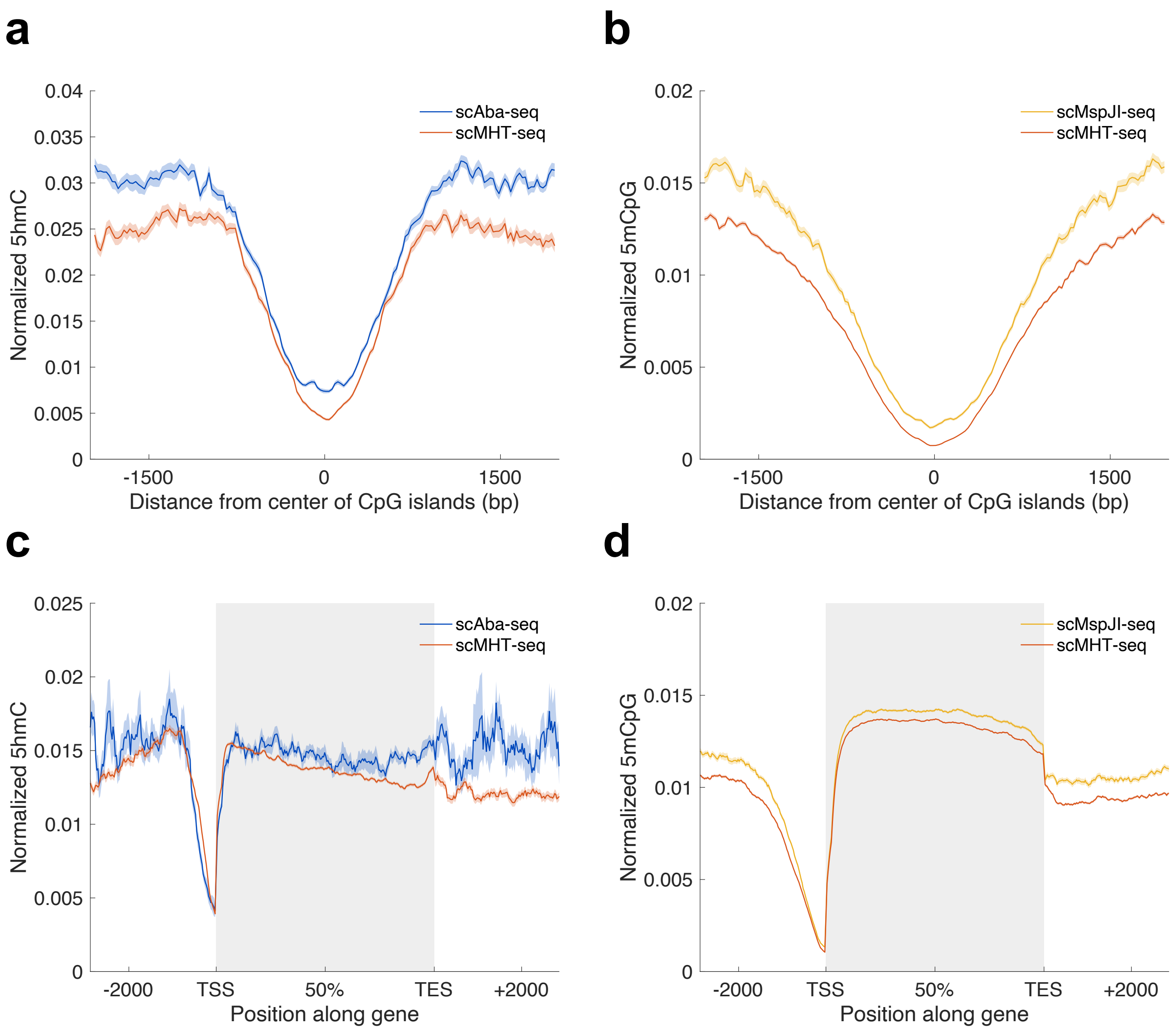


**Supplementary Figure 5 | Genome-wide 5mC and 5hmC landscapes at CpG islands and genes.** (**a,c**) 5hmC profiles at CpG islands and genes obtained using scAba-seq and scMHT-seq. The genome-wide profiles show low levels of 5hmC at CpG islands and at gene promoters. (**b,d**) 5mCpG profiles at CpG islands and genes obtained using scMspJI-seq and scMHT-seq. The genome-wide profiles show hypomethylation at CpG islands and at gene promoters. Solid lines indicate mean normalized reads over single cells and shaded regions indicate standard error of the mean over 10,000 bootstrap samples.

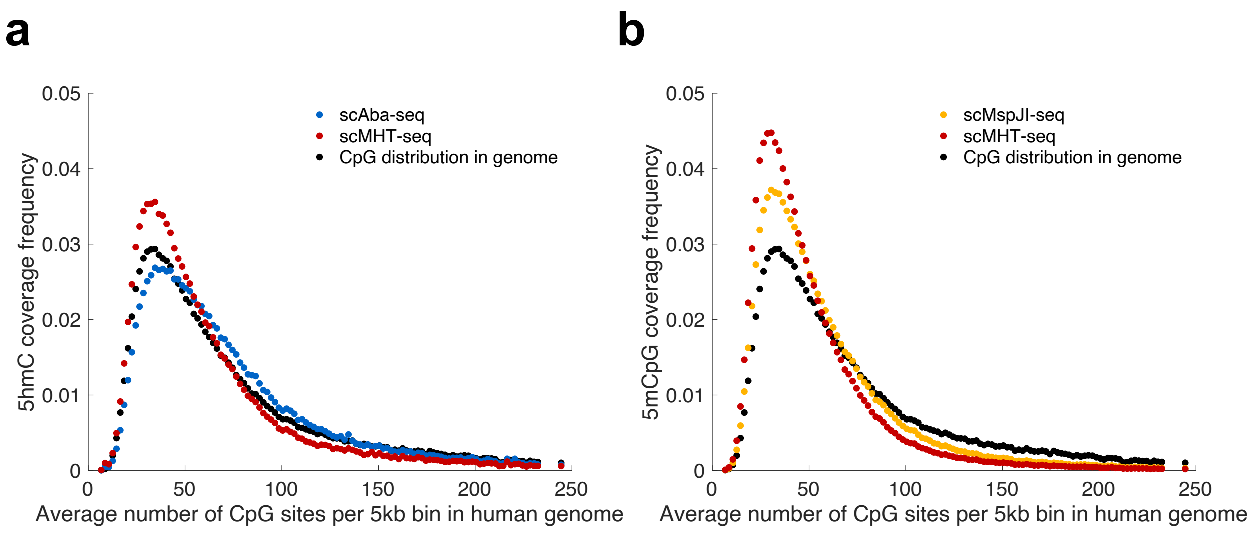


**Supplementary Figure 6 | Distribution of 5hmC and 5mCpG sites over genomic regions of varying CpG density.** (**a**) The distribution of 5hmC sites detected by scMHT-seq and scAba-seq over genomic regions of varying CpG density. (**b**) The distribution of 5mCpG sites detected by scMHT-seq and scMspJI-seq over genomic regions of varying CpG density. In both panels, the black curve shows the distribution of CpG sites over 5 kb bins in the human genome. The figure shows data from H9 hESCs.

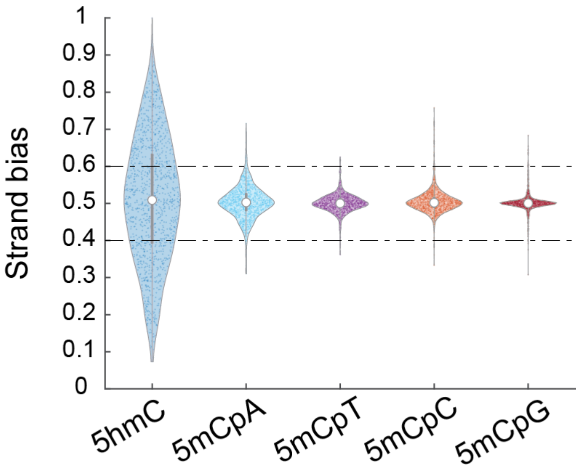


**Supplementary Figure 7 | 5hmC and 5mC strand bias in mouse embryonic stem cells.** Violin plots show chromosome-wide 5hmC and 5mC strand bias distributions in E14 mouse embryonic stem cells. Each point represents an individual chromosome in single cells. scAba-seq was used to quantify 5hmC strand bias using data from Mooijman *et al*[1]. scMspJI-seq was used to quantify 5mC strand bias in different dinucleotide contexts using data from Sen *et al*[2].

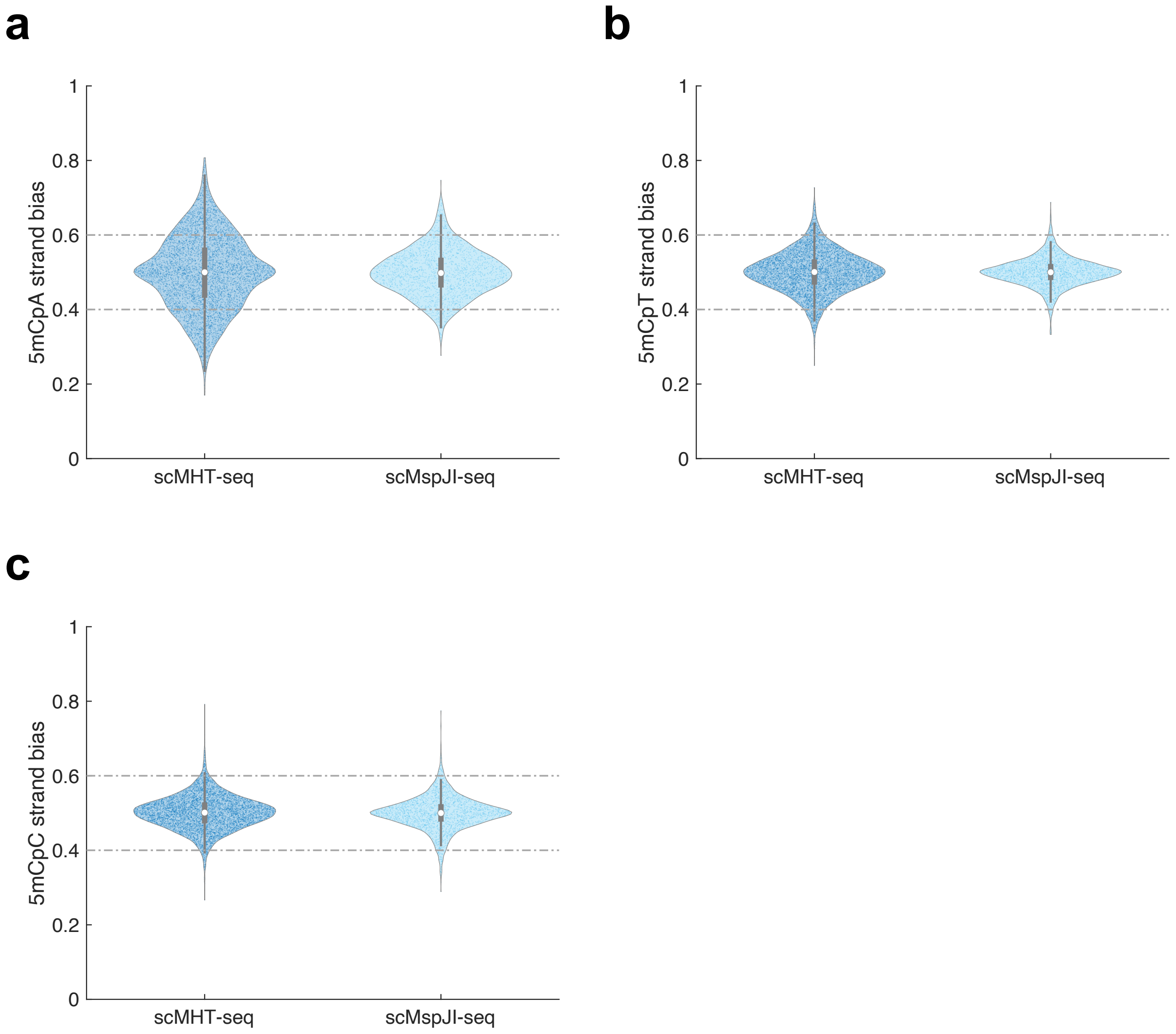


**Supplementary Figure 8 | Comparison of 5mCpH strand bias quantified using scMHT-seq and scMspJI-seq.** Violin plots show similar chromosome-wide 5mCpA, 5mCpT, and 5mCpC strand bias distributions obtained using data from scMHT-seq and scMspJI-seq. Each point represents an individual chromosome in single cells. These experiments were performed on H9 hESCs.

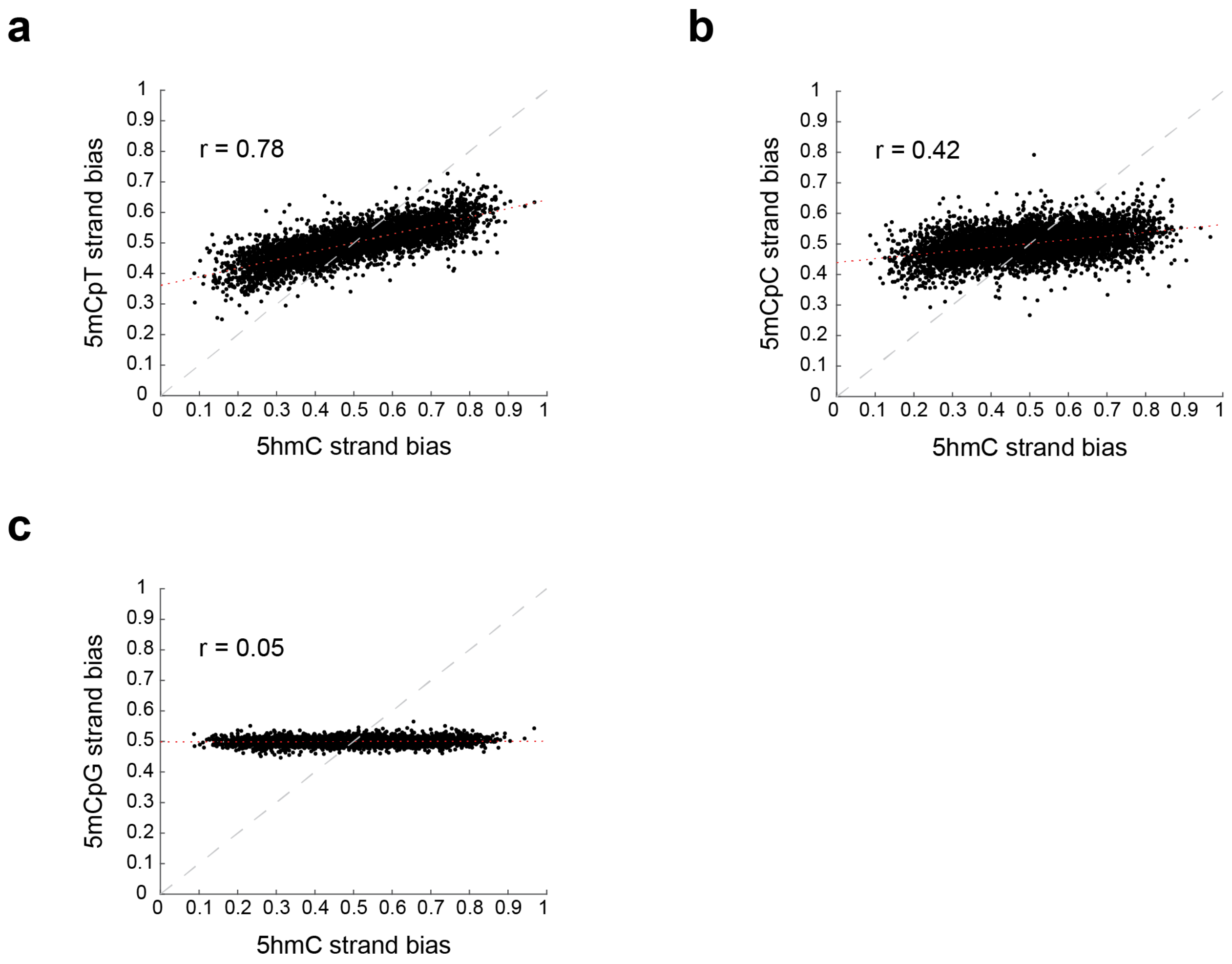


**Supplementary Figure 9. 5hmC and 5mCpH strand bias are highly correlated.** (**a-c**) Comparison of 5mCpT (a), 5mCpC (b), and 5mCpG (c) strand bias to 5hmC strand bias for individual chromosomes across all single H9 cells profiled by scMHT-seq. The best fit line is shown in dotted red. Pearson correlations are shown for each panel. These experiments were performed on H9 hESCs.

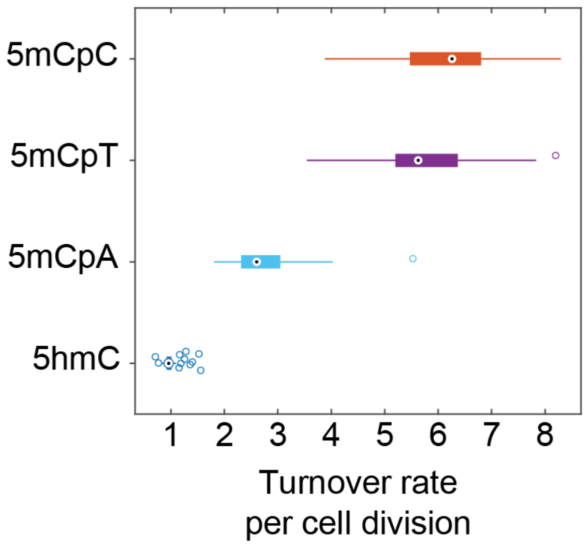


**Supplementary Figure 10 | Estimating the turnover rates of 5hmC and non-CpG methylation per cell division.** We fit experimental strand bias distributions using a stochastic model previously described by us in Mooijman *et al.* to estimate the net turnover rate per cell division of a DNA modification on a new DNA strand relative to the older DNA strand on a chromosome[1]. Boxplots show the results from 100 independent stochastic simulations. These turnover rates are estimated for H9 hESCs.

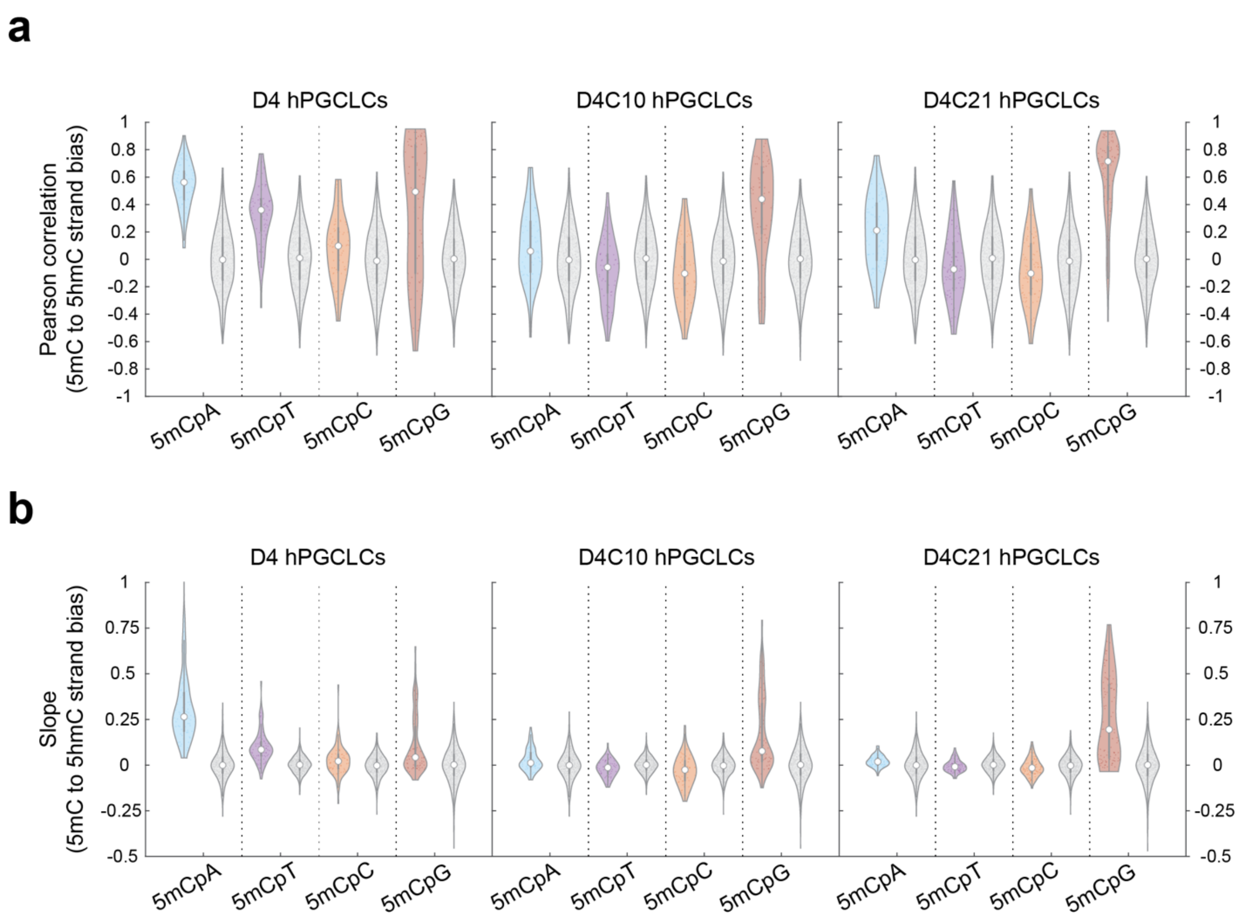


**Supplementary Figure 11 | Comparison of 5hmC stand bias to 5mC strand bias in hPGCLCs.** (**a,b**) The violin plots compare 5hmC strand bias to 5mC strand bias for the same chromosome in individual hPGCLCs derived from different time points in extended culture. Each point indicates the Pearson correlation (a) or slope (b) of a single cell. Gray dots within the gray violin plots indicate an *in silico* cell, where strand bias of each DNA modification on a chromosome was randomly sampled from the experimental dataset.

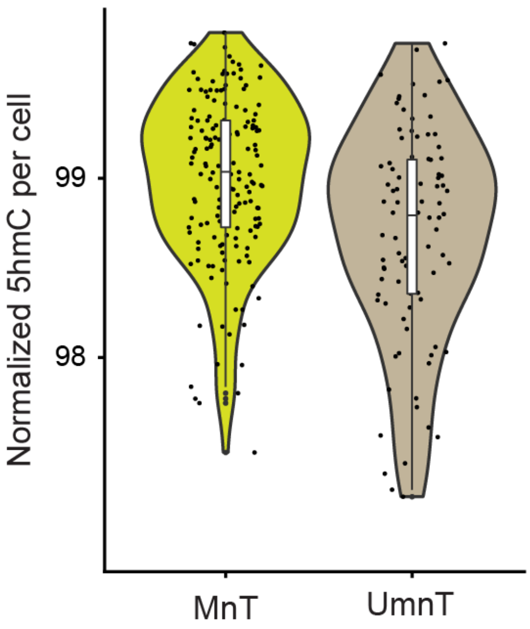


**Supplementary Figure 12 | Normalized 5hmC levels in MnT and UmnT hPGCLCs.** Violin plot shows the distribution of normalized 5hmC levels in individual hPGCLCs that display high (MnT) or impaired (UmnT) DNMT1-mediated maintenance methylation. Normalized 5hmC levels are obtained by taking the ratio of endogenous 5hmC sites detected in individual cells relative to 5hmC spike-in molecules added to each reaction well containing a single cell.

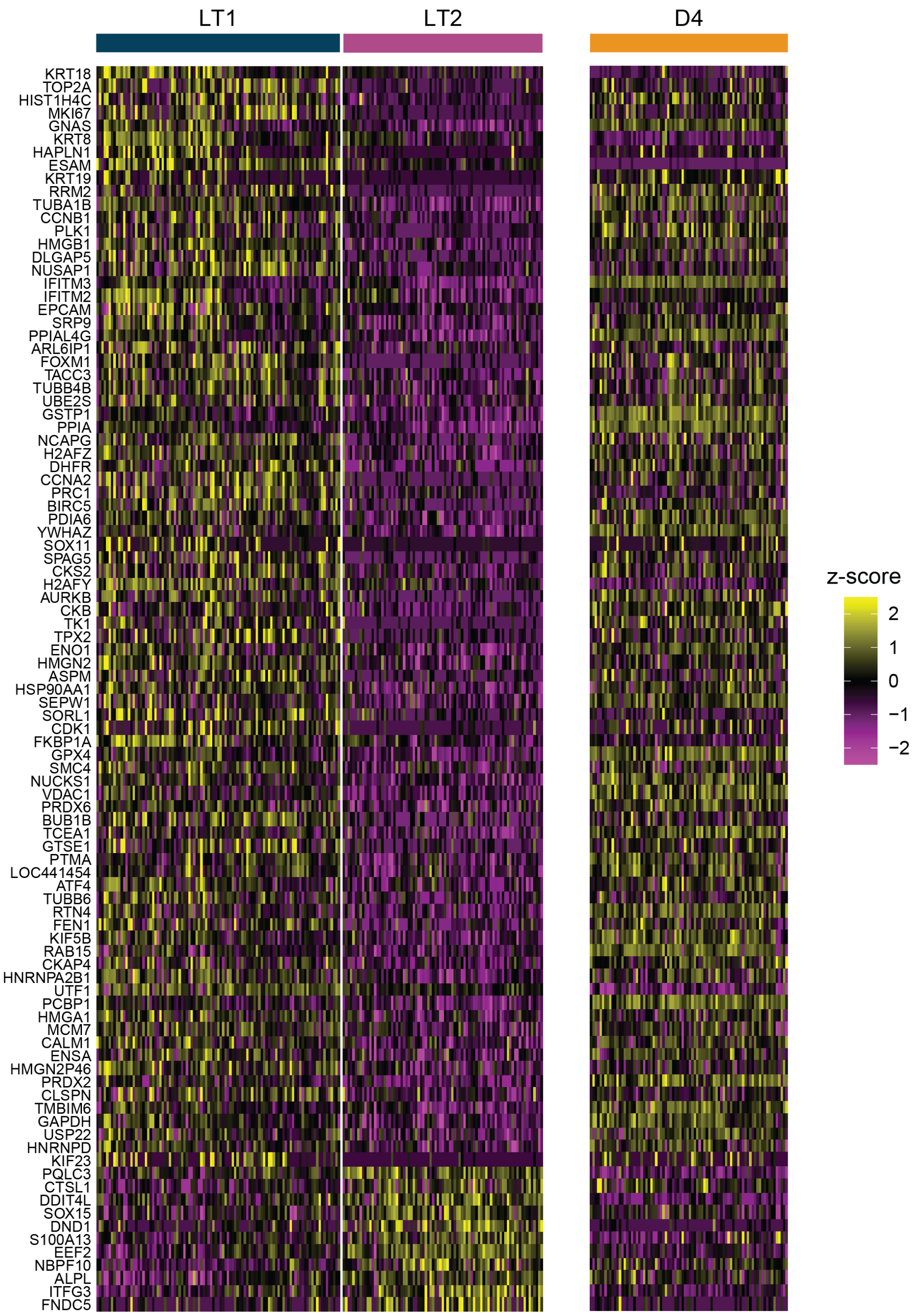


**Supplementary Figure 13 | Gene expression differences between hPGCLCs in the LT1 and LT2 transcriptional groups.** Heatmap of differentially expressed genes between the two hPGCLC populations in long term culture, LT1 and LT2. The corresponding expression in the D4

transcriptional group is also shown. Color indicates z-score of normalized gene expression across all hPGCLCs.
